# Inferring Synaptic Excitation/Inhibition Balance from Field Potentials

**DOI:** 10.1101/081125

**Authors:** Richard D. Gao, Erik J. Peterson, Bradley Voytek

## Abstract

Neural circuits sit in a dynamic balance between excitation (E) and inhibition (I). Fluctuations in this E:I balance have been shown to influence neural computation, working memory, and information processing. While more drastic shifts and aberrant E:I patterns are implicated in numerous neurological and psychiatric disorders, current methods for measuring E:I dynamics require invasive procedures that are difficult to perform in behaving animals, and nearly impossible in humans. This has limited the ability to examine the full impact that E:I shifts have in neural computation and disease. In this study, we develop a computational model to show that E:I ratio can be estimated from the power law exponent (slope) of the electrophysiological power spectrum, and validate this relationship using previously published datasets from two species (rat local field potential and macaque electrocorticography). This simple method--one that can be applied retrospectively to existing data--removes a major hurdle in understanding a currently difficult to measure, yet fundamental, aspect of neural computation.

## INTRODUCTION

Neurons are constantly bombarded with spontaneous synaptic inputs. This state of fluctuating activity is often referred to as the high-conductance state (Destexhe et al. 2003), and gives rise to the asynchronous, irregular (Poisson-like) firing observed in-vivo (Destexhe et al. 2001). In this state, neural circuits sit in a balance between synaptic excitation (E) and inhibition (I), typically consisting of fast glutamate (AMPA) and slower GABA (GABA_A_) inputs, respectively, where inhibitory conductance is typically two to six times the strength of excitatory conductance (Xue et al. 2014; Alvarez & Destexhe 2004). Physiologically, the balanced push-pull of E:I interaction is essential for neuronal homeostasis (Turrigiano & Nelson 2004) and the formation of neural oscillations (Atallah & Scanziani 2009). Computationally, E:I balance allows for efficient information transmission and information gating (Vogels & Abbott 2009; Salinas & Sejnowski 2001), network computation (Mariño et al. 2005), and working memory maintenance (Lim & Goldman 2013). Conversely, an imbalance between excitation and inhibition, where one dominates the other either during key developmental periods or tonically thereafter, is implicated in neurological and psychiatric disorders such as epilepsy (Symonds 1959; González-Ramírez et al. 2015), schizophrenia (Uhlhaas & Singer 2010), and autism/Rett Syndrome (Dani et al. 2005; Mariani et al. 2015; Rubenstein & Merzenich 2003), as well as impairments in information processing and social exploration (Yizhar et al. 2011).

Given such a state of intricate balance and its profound consequences when disturbed, quantifying the E:I ratio could aid in characterizing the functional state of the brain in better detail. Existing methods for estimating E:I ratio focus predominantly on interrogation of precisely selected cells, either through identification of excitatory and inhibitory neurons based on their extracellular action potential waveforms (Peyrache et al. 2012), or by intracellular voltage-clamp recordings to separately measure the depolarizing and hyperpolarizing synaptic currents (Monier et al. 2008), often combined with pharmacological or optogenetic manipulations (Xue et al. 2014; Reinhold et al. 2015). These methods are invasive and are restricted to a small population of cells, making them difficult to apply in clinical settings and to *in vivo* population-level analyses critical for understanding neural network functioning. Other methods, such as magnetic resonance spectroscopy (Henry et al. 2011) and dynamic causal modeling (DCM) (Legon et al. 2015), are able to provide much greater spatial coverage, thus enabling the sampling of E:I ratio across the brain. However, this gain in coverage comes at a cost of temporal resolution – requiring several minutes of data for a single snapshot – and are based on restrictive connectivity assumptions.

Here, we aim to address this important gap in methodology to measure E:I ratio with broad population coverage and fine temporal resolution. Two recent lines of modeling work motivate our hypothesis. First, it has been shown that synaptic input fluctuations during the high conductance state can be accurately modeled by a summation of two stationary stochastic processes representing excitatory and inhibitory inputs (Alvarez & Destexhe 2004). These inputs have different rates of decay, corresponding to a faster AMPA current and a slower GABA_A_ current, which can be readily differentiated in the frequency domain and computationally inferred from single membrane voltage traces (Pospischil et al. 2009). Second, population-level neural field recordings, such as the local field potential (LFP) and electrocorticography (ECoG), have been shown to be primarily dominated by postsynaptic currents (PSC) across large populations (Miller et al. 2009; Mazzoni et al. 2015; Buzsáki et al. 2012), and recent work by (Haider et al. 2016) observed tight coupling between the LFP and both excitatory and inhibitory synaptic inputs in the time domain. Thus, we combine these two findings and reason that changes in the relative contribution between excitatory and inhibitory synaptic currents must also be reflected in the field potential, and in particular, in the frequency domain representation (power spectral density, or PSD) of LFP and ECoG recordings. In this work, we derive a straightforward metric that closely tracks E:I ratio via computational modeling, and demonstrate its empirical validity by reanalyzing publically available databases from two different mammalian species.

## RESULTS

### E:I ratio drives 1/f changes in simulation

To model LFP under the high conductance state, we simulate an efferent “LFP” population receiving independent Poissonic spike trains from an excitatory and an inhibitory population (Figure 1A). Excitatory inputs depolarize target neurons at AMPA synapses, which have a faster conductance profile compared to inhibitory GABA_A_ synapses (Figure 1B) (aggregate values from CNRGlab @ UWaterloo). Convolving the spike trains and their respective conductance profile in time, then weighting by the synaptic reversal potentials, results in integrated synaptic currents (Figure 1C, top), which are linearly summated to compute the population LFP (Mazzoni et al. 2015) (Figure 1C, bottom). In the frequency domain, we observe that the power spectral density of the LFP (LFP-PSD) follows a decaying (1/*f*) power law for frequencies past 20 Hz (negatively linear in log-log plot), which directly results from adding the two current components, both following power law decays (Figure 1D). Note that the current-PSDs begin decaying at different frequencies, due to the different rise and decay time constants of AMPA and GABA_A_ conductance profiles, which have been previously observed in intracellular models of the balanced, high conductance state (Destexhe & Rudolph 2004).

**Figure 1.**
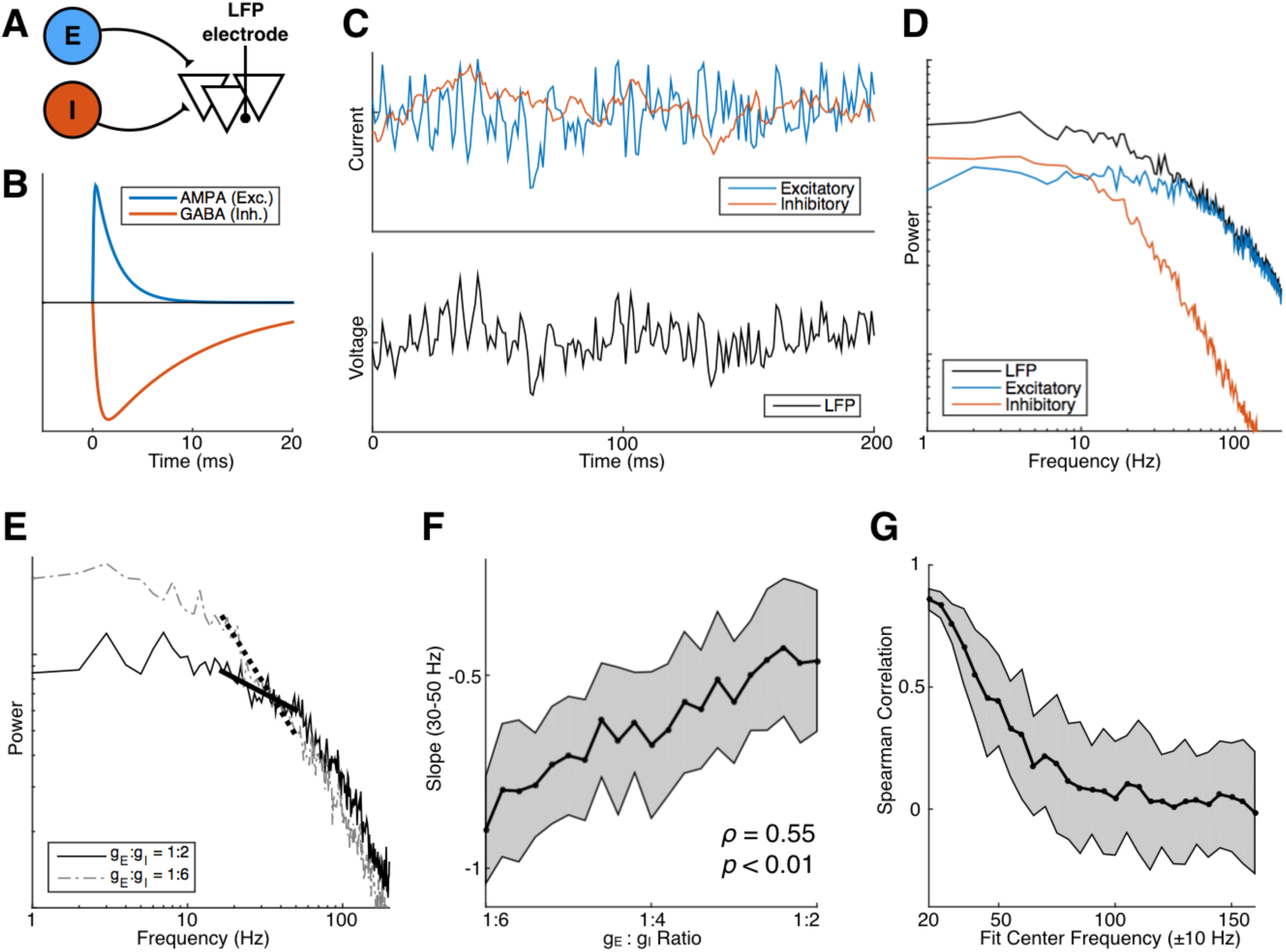
E:I ratio correlates with PSD slope in simulation. (A) Model schematic: an “LFP population” receives input from two Poisson populations, one excitatory and one inhibitory. (B) AMPA and GABA_A_ conductance profiles follow a difference-of-exponentials with different rise and decay time constants. (C) Example time trace of simulated total synaptic currents (top) and LFP (bottom). (D) PSDs of simulated signals in (C). Note power law decays in current-PSDs that begin at different frequencies. (E) Increasing E:I ratio from 1:6 to 1:2 causes a rotation, producing a flatter PSD. (F) E:I ratio is positively correlated with PSD slope between 30–50 Hz. (G) Positive rank correlations between E:I ratio and PSD slope diminish with increasing frequency of fitting window, up to 100 Hz.

By changing the relative contributions of excitation and inhibition (E:I ratio), we shift the frequency at which the current-PSDs cross over, which in turn produces different LFP-PSD slopes (power law exponent) in the intermediate frequency range (Figure 1E). To quantify this relationship, we vary E:I ratio from 1:2 to 1:6, and observe that LFP-PSD slope between 30 to 50 Hz positively correlates with E:I ratio (Figure 1F). The change in slope is restricted to only the low-to-intermediate frequency ranges (below 100 Hz), as we observe a steady decline in correlation between E:I ratio and PSD slope when slope is fitted across shifting, 20-Hz wide frequency windows (Figure 1G). For subsequent slope analyses, we use a 20-Hz window of the lowest possible frequencies that is above visible oscillatory peaks in the PSD, as a clear drop in correlation is observed when a narrowband oscillation, such as beta (15–25 Hz), is present (Figure S1). Additionally, we avoid high frequency regions because action potentials and firing rate changes have been shown to alter high gamma power at frequencies as low as 50 Hz (Manning et al. 2009; Ray & Maunsell 2011; Miller et al. 2007). In summary, our forward LFP model suggests that E:I ratio is monotonically related to LFP-PSD slope in a range between 30–70 Hz, when uncorrupted by oscillatory peaks, and that increasing E:I ratio increases (flattens) PSD slope.

### Depth-varying synapse density in rat CA1

To test the relationship between E:I ratio and PSD slope empirically, we first take advantage of the fact that excitatory and inhibitory synapse densities vary along the pyramidal dendrites in the CA1 region of the rat hippocampus (Megías et al. 2001). We ask: can changes in the ratio of excitatory to inhibitory synapse density be captured by changes in PSD slope, measured along the depth of CA1? Shank recordings are obtained from CRCNS data portal (Mizuseki et al. 2011; Mizuseki et al. 2009), sampling LFP at evenly spaced electrodes across a depth of 280 um centered (*post hoc*, see Methods) on the pyramidal cell layer in CA1 (Figure 2A). PSDs are computed using data from entire recording sessions of open field foraging (Figure 2B). PSD slopes are then fitted between 30–50 Hz to arrive at a slope profile that varied across depth (Figure 2C). To compute E:I ratio, we adapt synapse density values from (Megías et al. 2001) and spatially smooth it to produce data points at equivalent LFP electrode depths (Figure 2D and S2). We find that PSD slope across depth is significantly correlated with the AMPA to GABA_A_ synapse ratio (Spearman’s *ρ* = 0.23, *p* < 10^−5^), corroborating our *a priori* simulation results (Figure 2E). Interestingly, inhibitory synapse density alone correlates more strongly with PSD slope (Spearman’s *ρ* = −0.41, *p* < 10^−5^; Figure 2F). To further dissect the covariations, we create multivariate linear models regressing for slope, using every combination of excitatory density, inhibitory density, and E:I density ratio (Table S1). We find that each variable alone produces models that are significantly better than null (constant-only) and with coefficients in the direction expected (positive for E, E:I ratio; negative for I), where the full model with all 3 predictors achieves the highest adjusted *R*^2^. However, inhibitory density in any combination produces the largest increase in adjusted *R*^2^. Thus, we find that PSD slope significantly correlates with E:I ratio in the rat CA1, as measured by synapse density, though the effect is strongly driven by the presence of inhibition.

**Figure 2.**
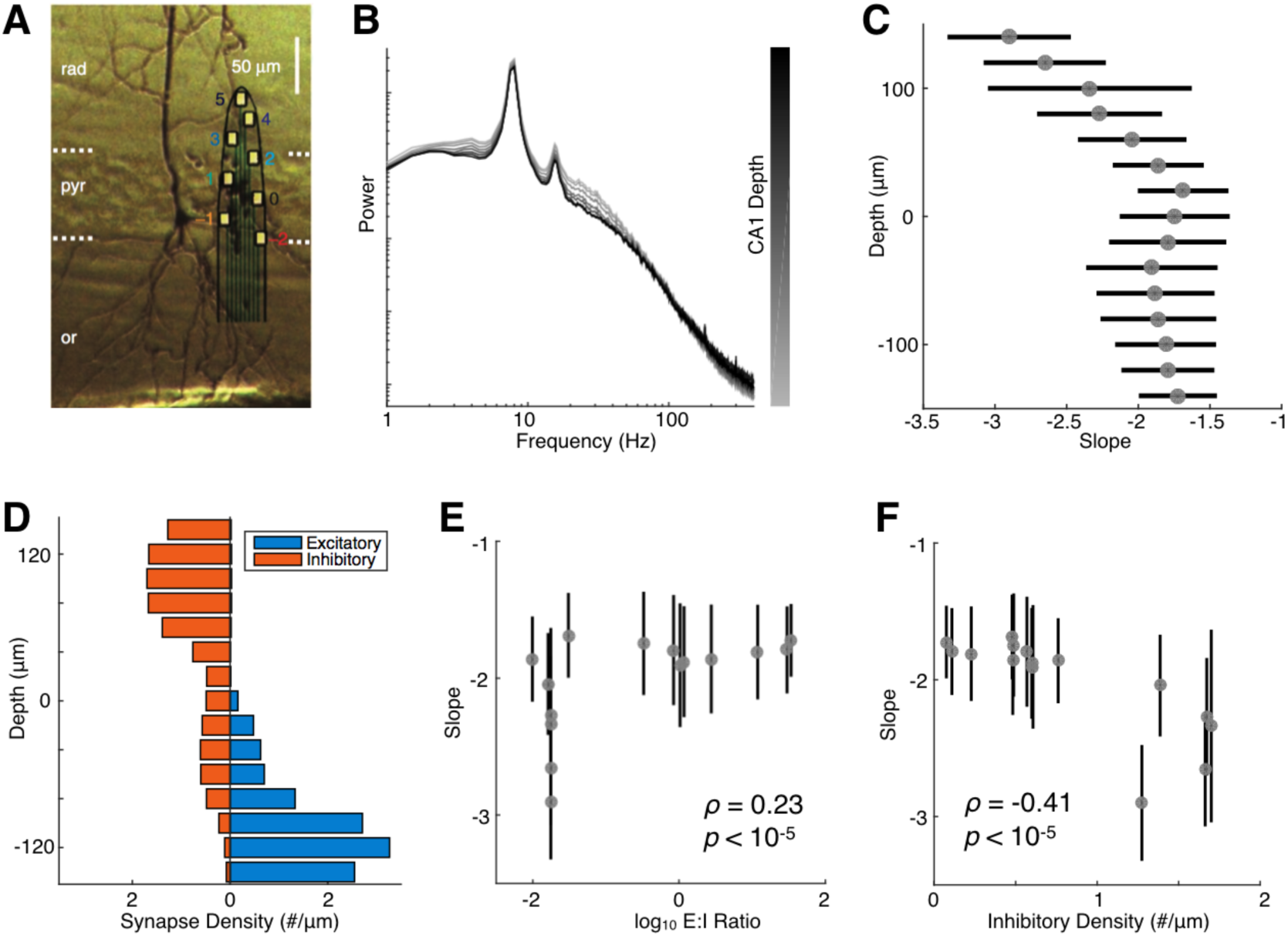
LFP-PSD slope varies with E:I synapse density ratio in rat CA1. (A) Example shank spanning across CA1 (*rad*: *stratum radiatum*; *pyr*: *stratum pyramidale*; *or*: *stratum oriens*; adapted from (Mizuseki et al. 2011)). (B) Example PSDs computed from electrodes along one recording shank. (C) Aggregate slope profile across depth, centered to the middle of pyramidal layer (0 μm) (horizontal bars denote standard deviation). (D) Excitatory (AMPA) and inhibitory (GABA_A_) synapse density varies across CA1 depth. (E and F) LFP-PSD slope correlates positively with E:I synapse density ratio (E) and negatively with GABA_A_ density (F) (vertical bars denote standard deviation).

### Theta-modulated cycles of excitation & inhibition

If LFP-PSD slope indeed tracks changes in the balance between excitation and inhibition, it should not only do so statically across space, but dynamically across time as well. Theta oscillation in the rat hippocampus reflects periodic bouts of excitation and inhibition (Buzsáki 2002). Therefore, we posit that PSD slope would be steeper during the inhibitory phase of theta, and flatter during the excitatory phase. To test this, we use the same CA1 dataset as above, and divide each LFP recording into temporal segments of peak and trough based on theta phase (Figure 3A). Fast Fourier Transforms (FFTs) are computed from these short segments and averaged, showing distinctive slope differences and enveloping the grand average PSD (Figure 3B).

**Figure 3.**
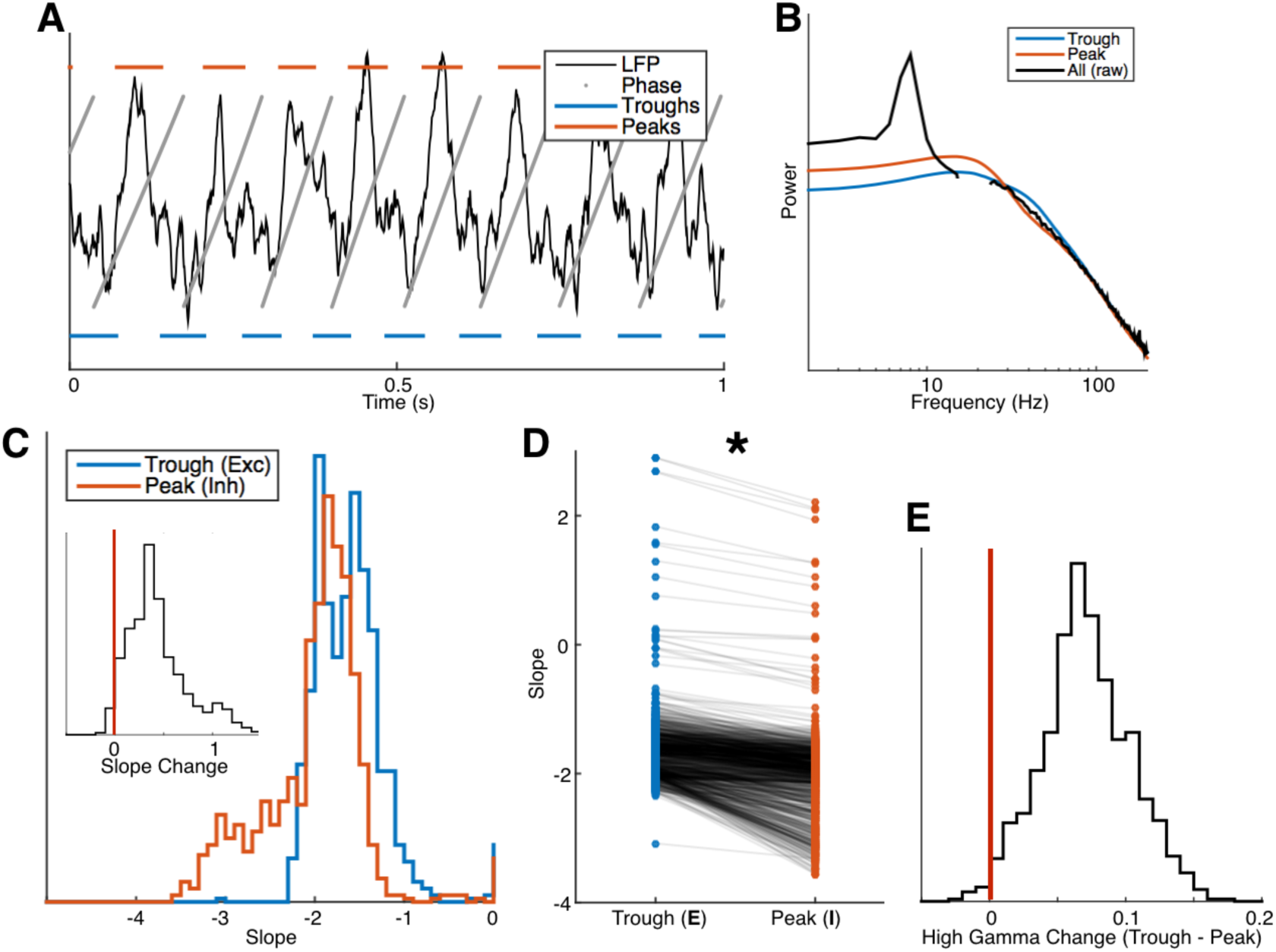
PSD slope tracks theta-modulated changes in E:I balance. (A) Schematic of how LFP segments are divided and binned based on theta phase. (B) Example PSD of a single channel over the entire recording (black, notch filter applied in beta range), and averages across all troughs (blue) and peaks (red) only. (C) Distribution of slope values shifts rightward (more positive) during theta troughs. Inset: distribution of difference in slope (trough minus peak) lies significantly above 0 (vertical red line). (D) Individual-channel comparison of slopes during theta troughs vs. peaks, each channel represented by a pair of connected dots showing nearly universally more negative slope during peaks compared to troughs (* *p* < 10^−5^). (E) Distribution of difference (trough minus peak) in high frequency activity (HFA, 140–230 Hz) in all channels lies significantly above 0 (vertical red line), indicating an increase in high gamma power from peak to trough.

We find that, across all channels, PSD slope (30–50 Hz) during theta peaks were significantly more negative (steeper) than during theta troughs (paired *t*-test, *p* < 10^−5^; Figure 3C and 3D). On a single channel basis, we fit linear slopes to each short segment FFT, and found 844 out of 946 channels with significantly flatter slopes during troughs (2-sample *t*-test, *p* < 10^−5^). From this we infer that theta troughs correspond to periods of excitation, which agrees with the biophysical view that negativity in the LFP is due to depolarization of membrane potential (Buzsáki et al. 2012). Additionally, we observe that high-frequency (140–230 Hz) power – a surrogate for spiking activity and ripples in the hippocampus (Schomburg et al. 2012) – is higher during theta troughs than peaks, further indicating the correspondence between LFP troughs and windows of excitation (Figure 3E). Taken together, we find evidence that PSD slope can dynamically track periods of excitation and inhibition facilitated by theta oscillations in the rat hippocampus.

### Propofol-induced increase in GABA_A_-mediated inhibition

Finally, having shown correlative evidence supporting the hypothesis, we aim to further test the simulation predictions through causal manipulations. Propofol is a general anesthetic that positively modulates the effect of GABA at GABA_A_ receptors (Concas et al. 1991), effectively decreasing the global E:I ratio. Thus, we query another openly available dataset (http://www.neurotycho.org), in which electrocorticogram (ECoG) from macaque monkeys was recorded throughout sedation, to investigate whether ECoG-PSD slope reflects a decrease in E:I ratio induced via pharmacological manipulation (Yanagawa et al. 2013). PSDs are computed for all 128 recording channels per session, for awake resting and anesthetized conditions (Figure 4A). We observe a significant decrease in PSD slope after onset of anesthesia for all 4 recording sessions (paired *t*-test, all *p* < 10^−5^, Figure 4B). The slope decrease is strongest in frontal and temporal electrodes (Figure 4C), consistent with previous neuroimaging studies spatially locating propofol’s region of effect (Zhang et al. 2010). Interestingly, electrodes in the precuneus region show increases in PSD slope during anesthesia instead, suggesting a gain of activity, perhaps due to its situation as a critical, core node within the default mode network (Utevsky et al. 2014). Finally, to calculate temporally precise demarcations of consciousness state changes, we estimate PSD slope in a time-resolved fashion by fitting over 1-second long sliding FFTs across the entire recording session. We find that PSD slope dynamically tracks the stability of brain state during awake resting, followed by a rapid push towards inhibition after injection that is consistent with propofol’s time of onset (15–30 seconds), as well as the slow rebalancing during recovery from anesthesia (Figure 4D). Unexpectedly, we also observe a rapid increase in slope, back to resting-state values, following the initial gain in inhibition, suggesting a global re-normalization process. Overall, we demonstrate that ECoG-PSD slope dynamically tracks propofol-induced gain in inhibition consistently across brain regions and time.

**Figure 4.**
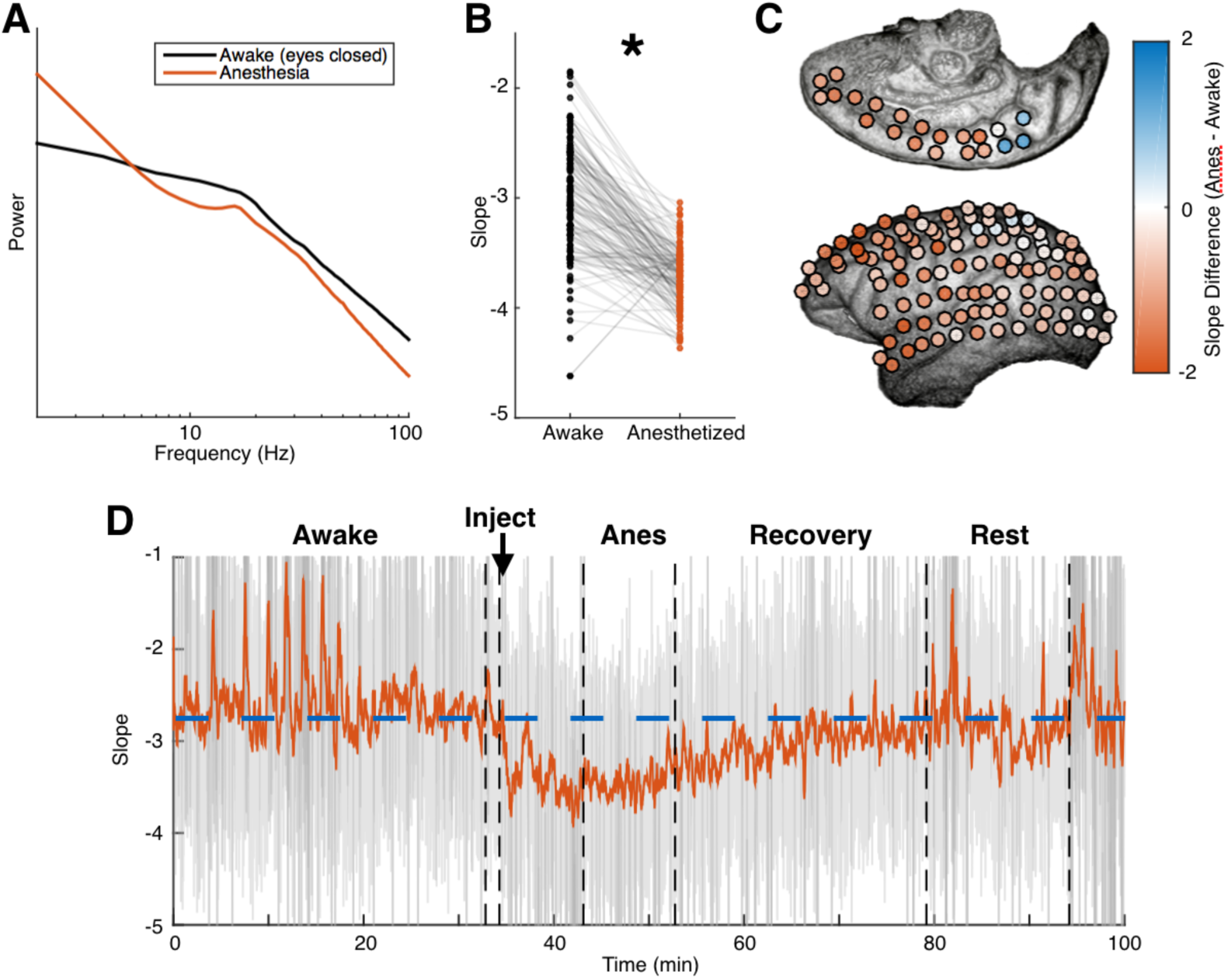
ECoG-PSD slope tracks propofol-induced global inhibition. (A) Average PSD across all channels during resting (black) and anesthetized (red) show distinct slope differences beyond 30 Hz. (B) Significant slope decrease is observed during anesthesia (pair *t*-test, * *p* < 10^−5^). (C) Slope decrease is observed across most of cortex, most prominently in the frontal and temporal areas. Slope increase is observed exclusively in the precuneus. (D) Time-resolved estimate of PSD slope tracks, with fine temporal resolution, changes in brain state from awake to anesthetized (Anes), and as well as a slow recovery to baseline rest levels (marked by dashed blue line). Grey, unsmoothed; red, 15s smoothing window applied.

## DISCUSSION

Guided by predictions from computational modeling, our analyses of existing datasets from two mammalian species with different experimental manipulations and recording equipment demonstrate that information about local E:I ratio can be robustly captured from the spectral representation of electrophysiological signals. Specifically, we show that LFP-PSD slope correlates with both anatomical E:I ratio--represented by changes in synaptic density ratio across CA1 layers--and dynamic E:I ratio as modulated by theta oscillation in the rat hippocampus. In addition, ECoG-PSD slope tracks the increase of inhibition in non-human primate brains induced by propofol, across brain regions and time.

Evidence that spiking can be partially extracted from the broadband (2–250 Hz) or high gamma (>80 Hz) spectral power of meso-/macro-scale neural recordings (LFP, ECoG) provided an important link between local neuronal activity and the LFP, opening numerous avenues of research (Mukamel et al. 2005; Manning et al. 2009). In contrast to the copious literature regarding broadband/high gamma, much of the work on E:I balance has been limited to intracellular recordings, methods with limited temporal resolution, multiple single-unit recordings, or optogenetic manipulations. Given the broad and important role that E:I balance plays in neural computation, information transfer, and oscillatory and homeostatic mechanisms, the inability to measure E:I parameters easily and at a large scale has hindered basic and clinical research. To this end, we develop a simple metric that can be applied at different intracranial recording scales, which can potentially be extended to extracranial EEG recordings, with profound implications for clinical and basic science research.

### Limitations

There are several caveats in this study worth noting. Most notably is the underlying assumption that LFP and ECoG are solely composed of AMPA and GABA_A_ synaptic currents. In reality, LFP reflects the integration of all ionic currents, including action potentials (Schomburg et al. 2012) – which shift the broadband/high gamma frequencies (Manning et al. 2009; Mukamel et al. 2005; Miller et al. 2007) – and slow glial currents (Buzsáki et al. 2012). The computational model also makes several assumptions, such as homogeneous-rate spiking and constant PSC waveforms, as well as excluding biophysical details like 3D arrangement of the spiking population. These factors will certainly influence the overall shape of the PSD, although this class of LFP model we employ was shown to best approximate neuronal networks with 3D cellular morphology (Mazzoni et al. 2015), and has been used to capture the aforementioned broadband/high gamma relationship to spiking activity (Miller et al. 2009), a phenomenon that is also reproduced in our model through an overall (and equivalent) decrease in firing rate from both excitatory and inhibitory populations.

Finally, because non-neural sources such as the amplifier, reference scheme, and ambient noise can affect spectral slope, slope-inferred E:I ratio should only be interpreted in the context of a comparative experimental design in which the relative E:I ratio can be interrogated in response to experimental manipulations or population differences, rather than ascribing meaning to the exact value of the slope itself. In particular, it has been shown that different referencing schemes, such as bipolar vs. common-average, have profound effects on the measured PSD slope (Shirhatti et al. 2016). In addition, we observe that PSD slope of cortical ECoG is much more negative than that of CA1 LFP recordings, which, in turn, is lower than slopes produced by our LFP model, suggesting that anatomical differences and dendritic integration process all contribute to the measured slope (Lindén et al. 2010; Pettersen et al. 2014).

### Power Law (1/f) Decay in Neural Recordings

Power law exponent (slope) changes of the PSD (“rotation”) have recently been observed in several empirical studies, linking it to changes in global awake and sleep states (He et al. 2010), age-related working memory decline (Voytek et al. 2015; Voytek & Knight 2015), and visuomotor task-related activation (Podvalny et al. 2015). The *1/f* power law nature of neural recordings has been interpreted within a self-organized criticality framework (Bak et al. 1987; He et al. 2010), with general anesthesia argued to alter the criticality of self-organized brain networks (Alonso et al. 2014). It has been shown, however, that power law statistics do not imply criticality in neuronal networks (Touboul & Destexhe 2010), and the finding that neuronal activity exhibit power law statistics at all has been questioned (Bédard et al. 2006). Furthermore, many previous reports ignore or overlook the fact that PSD of neural recordings are not 1/f at all frequencies and do not have a constant power law exponent – both requirements in the self-organized criticality framework. Instead, LFP and ECoG PSDs often have constant spectral power at low frequencies between 1–10 Hz, as well as different power law exponents at different frequencies. For example, ultra-low frequency region (<1 Hz) was posited to exhibit 1/f decay due to recurrent network activity (Chaudhuri et al. 2016), and power law in the very high frequency (>200 Hz) was shown to be a result of stochastic fluctuations in ion channels (Diba et al. 2004).

Our model and results reconcile the 1/*f* and low-frequency plateau observation by the simple fact that the spectral representation of synaptic currents (Lorentzian) takes on that shape (Figure 1D), as others have noted before (Destexhe & Rudolph 2004). Furthermore, we propose that slope changes in a particular frequency region (30–70 Hz) correspond to changes in E:I balance, while making no claims about other frequency regions. Altogether, it follows that different processes may give rise to power law phenomenon at different temporal scales, hence different frequency ranges (Freeman & Zhai 2009). Our observations here do not negate the criticality perspective, but reframes it in E:I terms, wherein constant E:I balancing is crucial for maintaining neuronal excitability at a critical state (Xue et al. 2014).

In summary, our results overturn a long-standing challenge that the relative contributions of EPSCs and IPSCs to electrophysiological signals cannot be inferred (Yizhar et al. 2011). We show that this limitation can be overcome using relatively simple metrics derived from meso- and macro-scale neural recordings, and that it can be easily applied retrospectively to existing data, opening new domains of inquiry and allowing for reanalyses within an E:I framework. Furthermore, our results provide insights into several ongoing research domains, such as possible contributors to the *1/f* power law phenomenon often observed in field potential power spectra. By providing a new way for decoding the physiological information of the aggregate field potential, we can query brain states in novel ways, helping close the gap between cellular and cognitive neuroscience and increasing our ability to relate fundamental brain processes to behaviour and cognition as a result.

## EXPERIMENTAL PROCEDURES

*LFP simulation.* We simulate local field potentials under the high conductance state (Alvarez & Destexhe 2004), with the assumption that the LFP is a linear summation of total excitatory and inhibitory currents (Mazzoni et al. 2015). Poisson spike trains from one excitatory and one inhibitory population are generated by integrating interspike intervals (ISI) drawn from independent exponential distributions, with specified mean rate parameter. Each spike train is convolved with their respective conductance profiles, which are modeled as a difference-of-exponentials defined by the rise and decay time constants of AMPA and GABA_A_ receptors (Eq.1). The two resulting time series represent total excitatory (g_E_) and inhibitory (g_I_) conductances, respectively. E:I ratio is defined as the ratio of mean excitatory conductance to mean inhibitory conductance over the simulation time, and specific E:I ratios are achieved by multiplying the inhibitory conductance by a constant, such that mean g_I_ is 2–6 times mean g_E_. To calculate current, conductances are multiplied by the difference between resting potential (−65 mV) and AMPA and GABAA reversal potential, respectively. Local field potential (LFP), finally, is computed as the summation of the total excitatory and inhibitory current. All simulation parameters are specified in Table 1. PSD with oscillations is simulated by multiplying original 1/f PSD with oscillation masks in the frequency domain, amplifying power in the alpha (8–12 Hz) and beta (15–25 Hz) range. LFP power is normalized to unity for each E:I ratio.

Equation 1. Difference-of-exponential PSC in time domain

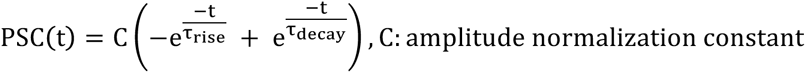

**Table 1.** LFP Simulation Parameters

| Parameter | Value |
| --- | --- |
| Population Firing Rate (E, I) | 2 Hz, 5 Hz |
| Population Size (E, I) | 8000, 2000 |
| Resting Membrane Potential | -65 mV |
| Reversal Potential (AMPA, GABA<sub>A</sub>) | 0 mV, -80 mV |
| Conductance Rise Time (AMPA, GABA<sub>A</sub>) | 0.1 ms, 0.5 ms |
| Conductance Decay Time (AMPA, GABA<sub>A</sub>) | 2 ms, 10 ms |
| E:I Ratio | 1:2 to 1:6 |

*Power spectral density (PSD)*. For all time series data (simulated and recorded LFP, ECoG), the PSD is estimated by computing the median of the square magnitude of the sliding window (short-time) Fourier transform (STFT). The median was used instead of the mean (Welch’s method) to account for the non-Gaussian distribution of spectral data, as well as to eliminate the contributions of extreme outliers. All STFT are computed with a window length of 1 second (2-seconds for CA1 data), and an overlap length of 0.25 seconds. A hamming window of corresponding length is applied prior to taking the FFT.

*1*/f *slope fitting*. To compute the 1/*f* power law exponent (*log-log* slope), we use robust linear regression (MATLAB *robustfit.m*) to find the slope and offset for the line of best fit over specified frequency ranges of the PSD (30–50 Hz, 40–60 Hz for monkey ECoG) (Eq.2).

Equation 2. Log-Log Linear Fit Parameter over Empirical PSD

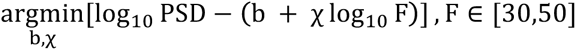

### Hippocampal LFP and CA1 depth analysis

LFP data (1250 Hz sampling rate) is recorded in stratum pyramidale of CA1 via 4 to 8 shank electrodes (200 um inter-shank distance), with 8 electrodes (160 um^2^ area) along the depth of each shank (20-um spacing), perpendicular to the pyramidal cell body layer (Mizuseki et al. 2009). PSD is computed for each electrode as specified above, and 1/f slope extracted. As in (Mizuseki et al. 2011), we align the shanks such that the electrode with the maximal ripple power (150 – 250 Hz) is set to position 0, the middle of stratum pyramidale. Other electrodes are vertically translated accordingly. This procedure iss repeated for all shanks in every recording (4 rats, 20 sessions total), resulting in slope estimates spanning a depth of 280 um, centered on the pyramidal layer. AMPA and GABA_A_ synapse densities are adapted from (Megías et al. 2001), for proximal stratum oriens and stratum radiatum dendrites, and smoothed with a 5-point Gaussian window to produce 15 data points at positions equivalent to LFP electrodes.

### Theta phase-modulated slope

Theta oscillation is first isolated with a FIR bandpass filter 5–12 Hz, (EEGLAB, *eegfilt.m*). Theta phase is computed as the complex phase angle of the Hilbert transform of the theta oscillation. Segments of theta phase are categorized as peak [−π/2 to π/2, through 0] or trough [π/2 to 3π/2, through π]. Each corresponding segment in the raw data (~75 samples) is then labeled as peak or trough, Hamming-windowed, and padded to 1250 samples. Average PSD for each phase category is computed as the median of all windowed FFT of the data segments of that category. 1/f slope is then fit to the average PSDs. Per-channel significance statistics are calculated by fitting 1/f slope to each individual cycle STFT for each channel and compared using two-sample *t*-test. To avoid power contamination in the short-time window estimates from observed beta oscillation, LFP data is notch filtered between 15–25Hz, see Figure S3 for filtered PSD. All results do not change when not filtered for beta, hence not presented above.

### Monkey ECoG During Anesthesia

ECoG data was collected from 2 macaque monkeys during rest, delivery of anesthesia (propofol, 5 & 5.2 mg/kg), and recovery (Yanagawa et al. 2013). PSD was computed for all ECoG channels (*n* = 128) for each experimental condition and fitted for 1/f slope. Due to clear gamma oscillation near 30 Hz biasing slope estimates, we fit over 40–60 Hz instead (Figure S4). We then compared slope fit differences at each electrode between conditions (paired-samples *t*-test). Time resolved slope fit was achieved by computing sliding window spectra (absolute value squared of FFT) throughout the duration of the recording (1 s window, 0.25 s step), and a slope estimate was computed for each window. A 15-second median filter was applied to smooth the slope time series plot for Figure 4D.

All simulation and analysis code can be found at www.voyteklab.com.

## ACKNOWLEDGEMENTS

We thank S.R. Cole, T. Donoghue, C. Holdgraf, R. van der Meij, E. Mukamel, D. Nitz, T. Noto, J. Olson, B. Postle, and T. Tran for invaluable discussion and comments, the Buzsáki Lab and CRCNS for their public repository of rat LFP data, and the Fujii Lab and NeuroTycho for their public repository of monkey ECoG data. B.V. is supported by the University of California, San Diego, Qualcomm Institute, California Institute for Telecommunications and Information Technology, Strategic Research Opportunities Program, and a Sloan Research Fellowship. R.D.G is supported by NSERC PGS-D, the Katzin Prize, and Frontiers of Innovation Scholars Program at the University of California, San Diego. The authors declare no conflicts of interest.

## AUTHOR CONTRIBUTIONS

R.D.G., E.J.P., and B.V. initiated and designed the study. R.D.G., E.J.P., and B.V. developed the computational model. R.D.G. analyzed the data. All authors discussed the results, while R.D.G., E.J.P., and B.V. wrote the manuscript.

---

**Figure S1.**
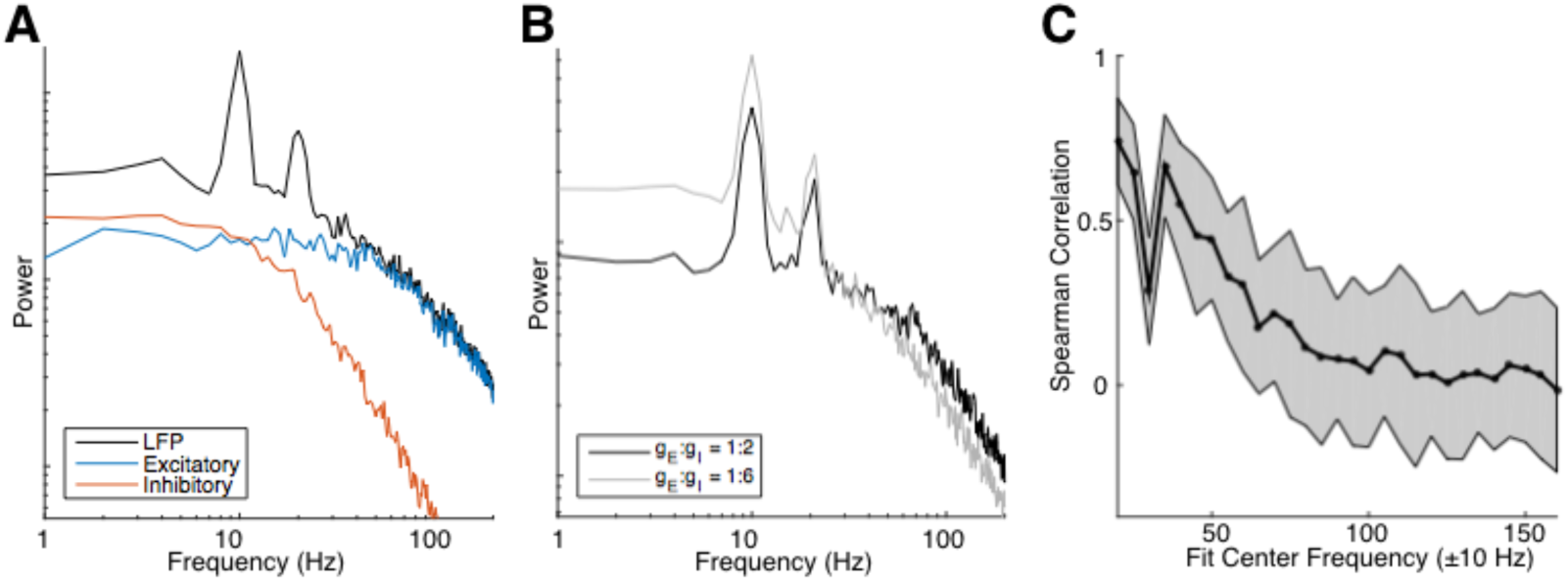
Simulated LFP with embedded oscillation corrupts 1/f fitting. (A) LFP- and current-PSDs with the addition of alpha and beta oscillations. (B) Changing E:I ratio produces slope changes in LFP-PSD with oscillations. (C) Oscillatory bump in the PSD corrupts 1/f fit, as seen by the drop in correlation coefficient when fitting window is centered on 30 Hz.

**Figure S2.**
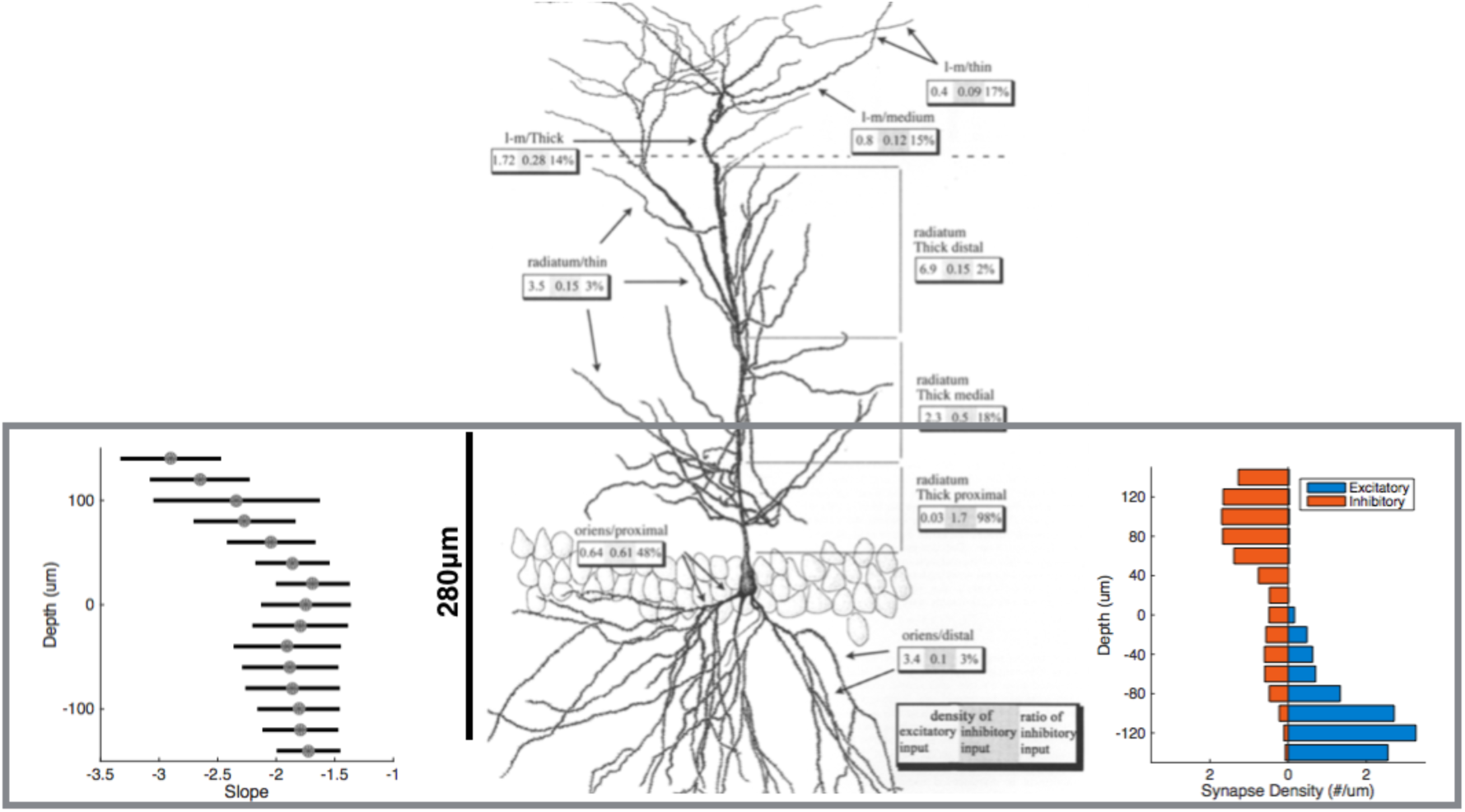
Region of interest in CA1. Synaptic density values corresponding to the segment of CA1 LFP data were taken from Megias et al, 2001. Figure 2 shows aligned slope and E:I density variations along the depth of interest, centered on the middle of the pyramidal layer, highlighted by the gray box. Cell morphology figure taken directly from Megias et al, 2001.

**Figure S3.**
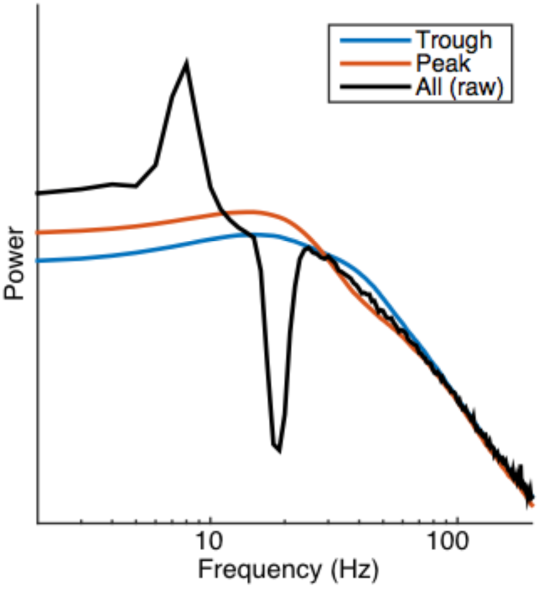
PSD with notched beta frequency, corresponding to Figure 3B in main text. Both figures are identical, with only the filtered region removed for the main text for aesthetic reasons.

**Figure S4.**
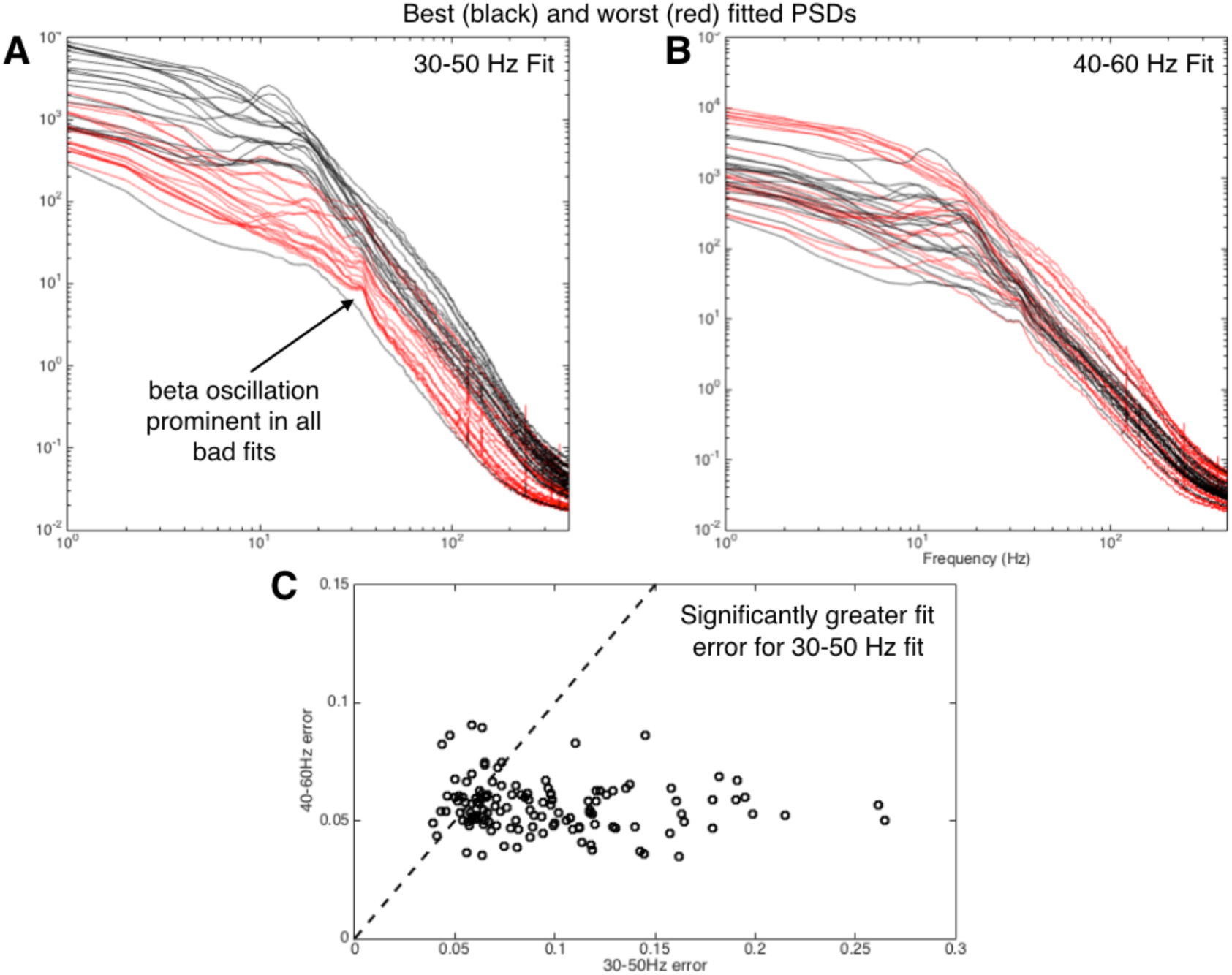
Fitting error for monkey ECoG. (A and B) 10 PSDs of best (black) and worst (red) fit, corresponding to lowest and highest mean squared error from robust fit. (A) shows fitting in the 30–50 Hz region, showing all channels of worst fits with prominent beta oscillations, whereas the problem is avoided when fit over the more linear regions of 40–60 Hz (B). (C) Fit errors for 30–50 Hz fit are significantly greater than when fit over 40–60 Hz for majority of channels, leading to the analysis decision of choosing the latter for analysis.

**Table S1.**
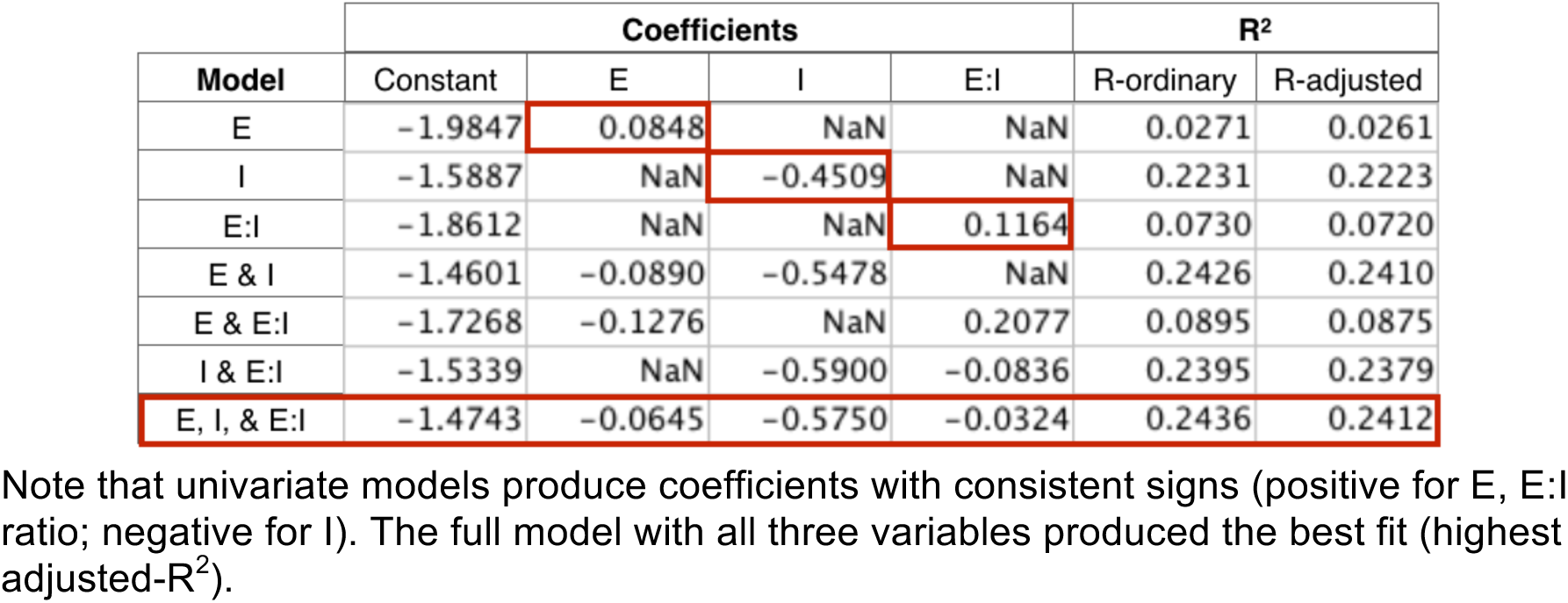
Multivariate Linear Model Coefficients and R^2^ for Slope vs. E, I, and E:I Ratio

